# Isolation of a novel *Sphingomonas* strain able to degrade the pleuromutilin tiamulin: omic analysis reveals its transformation pathway

**DOI:** 10.1101/2024.12.16.628731

**Authors:** Chiara Perruchon, Niki Tagkalidou, Natasa Kalogiouri, Eleni Katsivelou, Panagiotis A. Karas, Urania Menkissoglu-Spiroudi, Sotirios Vasileiadis, Dimitrios G. Karpouzas

## Abstract

Tiamulin (TIA) is a commonly used veterinary antibiotic, persistent in the animal digestive system and downstream receiving environments like soil after manuring. We aimed to isolate and characterize TIA-degrading bacteria for bioaugmentation strategies towards mitigating TIA environmental pressure. A strain able to degrade TIA and use it as sole C source was isolated from a soil exhibiting enhanced biodegradation of the antibiotic. The isolate degraded TIA at concentrations up to 100 µg ml^-1^, with pH and temperature optima of 6.5-7.5 and 16-25°C respectively. Phylogenomic analysis deemed the isolate to be a new *Sphingomonas* species (83.87 % ANI with *S. laterariae*, ≤ 95 %), which was named *Candidatus* Sphingomonas perruchonii. Genomics and transcriptomics revealed the TIA-driven induction of features conducive with its antiotrophic character; genes encoding for drug efflux pumps and the catabolism of xenobiotics. These included ribosome protective ABC-F transporters and efflux pumps able to protect the ribosome and microbial cells from TIA, oxygenases (e.g. P450 cytochrome) and hydrolases (e.g. alpha/beta hydrolases and amidohydrolases) possibly contributing to its degradation. LC-MS/MS analysis detected putative transformation products (TPs) of TIA, leading us to propose a transformation pathway. This involved a primary oxidation of the tricyclic moiety of TIA to a mono-hydroxylated derivative (TIA-O), potentially mediated by the highly upregulated monoxygenases, which was either further oxidized to TIA-O2 or hydrolysed, by the upregulated hydrolase or amidohydrolases, to 2-diethylamino-ethyl-thio acetic acid, both not degrading further. Further tests and in vitro functional analysis will verify the role of these genes in the transformation of TIA.

**IMPORTANCE:** Anthropogenic antibiotic pressure in environmental settings has been acknowledged as a major threat for the public health and the environment, through the: (i) associated ecotoxicological impact; and (ii) the increase of antibiotic resistance dispersal among non-pathogenic and pathogenic bacteria, restricting available choices for treating infections in health care. *In situ* microbial biotransformation of antibiotics and eventual removal is an environmentally friendly method currently under intense investigation, which was recently proposed for the reduction of antibiotic selective pressure. The present study provides mechanistic insights on the genomic constituents of antibiotrophy, namely, the tolerance against and the biodegradation/transformation of the antibiotic in question. We characterize the capacity of a bacterial isolate to transform tiamulin, a heavily used antibiotic in pig farming, and propose associated biotransformation mechanisms, through an integrated approach of standard microbiological methods combined with and multi-omics (genomics, transcriptomics, metabolomics).

## INTRODUCTION

Tiamulin (TIA) is a pleuromutilin antibiotic used exclusively in veterinary medicine to prevent and treat different enteric and respiratory diseases in animals, mainly pigs and poultry (1, 2). Previous studies have demonstrated the presence of TIA in pig and poultry faeces and liquid manure (3, 4), as well as in swine wastewaters (5). After administration, most of the applied dose is excreted through urine (35% for pigs) and animal faeces (65% for pigs) (1). C. Perruchon et al. (6) showed that TIA persisted in faeces stockpiled under ambient conditions (DT_90_ values ranging from 19.3 to 119.3 days) and also during anaerobic digestion (DT_90_ >365 days) exhibiting dose-dependent dissipation motifs. The high persistence of TIA in liquid pig manure under anaerobic storage conditions was also reported by M. P. Schlüsener et al. (7) with negligible degradation observed after 180 days.

It is, therefore, evident that the common practice of using faeces as manures for the fertilization of agricultural soils (8–10) involves a high risk of dispersal of TIA residues in soil (11), with possible adverse effects on the environment, the human and animal health. These could be manifested as: (a) contamination of natural water resources (12); (b) toxic effect on the soil microbial community, affecting soil quality and functioning (13); and (c) dispersal of antibiotic resistance genes (ARGs) (14, 15) that can be further transmitted to non-agricultural settings and the general public (16–18). Specific and efficient mitigation measures are necessary to minimize the risk of carryover of TIA residues in agricultural soils. Considering the lack of efficient alternative treatment methods, as mentioned above for anaerobic digestion and stockpiling (6), bioremediation could be a valuable tool for the treatment of TIA contaminated manures before their application in the agricultural environment. The basis of the development of such a strategy is the availability of an antibiothroph, a microorganism able to use an antibiotic, in this case TIA, as a nutrient source (19, 20).

Antibiotrophy relies primarily on the capacity of the microorganism to resist the biocidal action of the antibiotic and secondly on the evolution of catabolic traits that will enable the use of the antibiotic as energy source (19, 20). The increasing global consumption of antibiotics in the last decade in both human but mainly in veterinary medicine has led to the long-term exposure of soil microbial communities to antibiotics (21–23). This progressively resulted in an adaptive response of the soil microbiota through the development or enrichment and spread of ARGs, and the evolution of antibiotic-degrading functions (21–23). The role of soil microorganisms in the degradation of antibiotics was demonstrated in several studies where the DT_50_ values of antibiotics were significantly higher in sterilized *vs* non-sterilized soil samples (24–26). Furthermore, a growing body of literature reports the isolation of veterinary antibiotic-degrading microorganisms (27). E. Topp et al. (28) isolated a sulfamethazine-degrading *Microbacterium* sp. from an agricultural soil following long-term exposure. X. Yang et al. (29) isolated two penicillin-degrading *Bacillus* strains from soil collected from a pig farm, while bacterial strains belonging to *Stenotrophomonas* (30) and *Arthrobacter* (31) were capable of degrading tetracycline.

Little is known regarding the biodegradation of TIA in soil. T. K. X. Nguyen et al. (32) isolated three fungi from a swine farm that were able of TIA removal through the possible involvement of manganese peroxidase. More recently the same group reported the isolation of four bacterial consortia from swine wastewaters able to degrade TIA at concentrations of 2.5–200 mg/L with efficiencies of 60.1–99.9% over 16 h (33). The main members of the consortia were identified but neither the delineation of the roles of the members of the enriched consortia, nor the TIA transformation pathway elucidation, were further pursued. In the present study we aimed to isolate and characterize at genomic, functional and metabolic level a TIA antibiotrophic microorganism with the long-term aim to use it in bioaugmentation strategies for curation of animal manures prior to their application in agricultural soils. In this context, a novel soil bacterial species was isolated via enrichment culture from a soil where accelerated dissipation of TIA was previously observed (34). Its capacity to degrade TIA was evaluated in a series of liquid culture experiments. Genomics, transcriptomics and non-target metabolomics provided insights into the transformation pathway of TIA and identified genes with potential role in the transformation of TIA by the isolated microorganism.

## RESULTS

### Isolation of a TIA-degrading bacterium and characterization of its degrading capacity

A soil with demonstrated accelerated TIA dissipation (6) served as source for the TIA-degrading isolates.

### Isolation and initial taxonomic classification

The enriched soil was inoculated in two minimal media, mineral salt medium plus casamino acids (0.15 g l^-1^) which was either supplemented (MSMN) or not supplemented (MSM) with nitrogen (35) with TIA being the sole carbon source. Complete degradation of TIA occurred within 7 days, even during the first cycle of the liquid enrichment (Figure S1). Further acceleration in the degradation rates was observed from cycle to cycle, with complete disappearance of TIA occurring within 3 days in the final (fourth) enrichment cycle. Several cycles of plate-spreading and re-culturing (without/with antibiotics) led to the isolation of a culture that was able to rapidly degrade TIA in the presence of ampicillin (5 µg ml^-1^) (Figure S2). Further plating recovered two bacterial colonies, named 7 and 8, which upon selection, cultivation and verification of their degrading ability in liquid cultures showed equivalent TIA transformation ability in both MSM and MSMN (Figure S2).

The full length 16S rRNA gene sequences of the bacteria in the two cultures were identical and showed high 16S rRNA gene identity with the corresponding gene of *Sphingomonas laterariae* after search of the non-redundant database of the National Center for Biotechnology information (NCBI) using the basic local alignment search tool (BLAST; a more robust phylogenomics classification is provided further on). The 100% identity among the 16S rRNA gene sequences led to the choice of one of them for all subsequent tests.

### Effects of TIA concentration, pH and temperature on the degrading capacity and growth dynamics of the isolate

Complete TIA dissipation occurred within 1, 2 and 3 days when tested at nominal concentrations of 10, 50 and 100 µg ml^-1^ respectively (Figure 1A). No appreciable dissipation of TIA was observed at concentrations of 250 mg L^-1^ and above (Figure 1A) and in the non-inoculated controls (Figure S3). Regarding pH, TIA dissipation was evident at all tested pH levels, with a significant delay at pH 4.5 and maximum dissipation rates at pH 6.5 and 7.5 (Figure 1B; optimal parabola modelled pH of 7.14, Figure S4A). Finally, out of the tested temperatures (4, 16, 25, and 37 °C), the most rapid degradation of TIA was evident at 25°C and 16°C (Figure 1C), while only partial or no degradation was evident at 4°C and 37°C respectively (Figure 1C; optimal 3^rd^ order polynomial modelled temperature 20.9 °C, Figure S4B).

**Figure 1.**
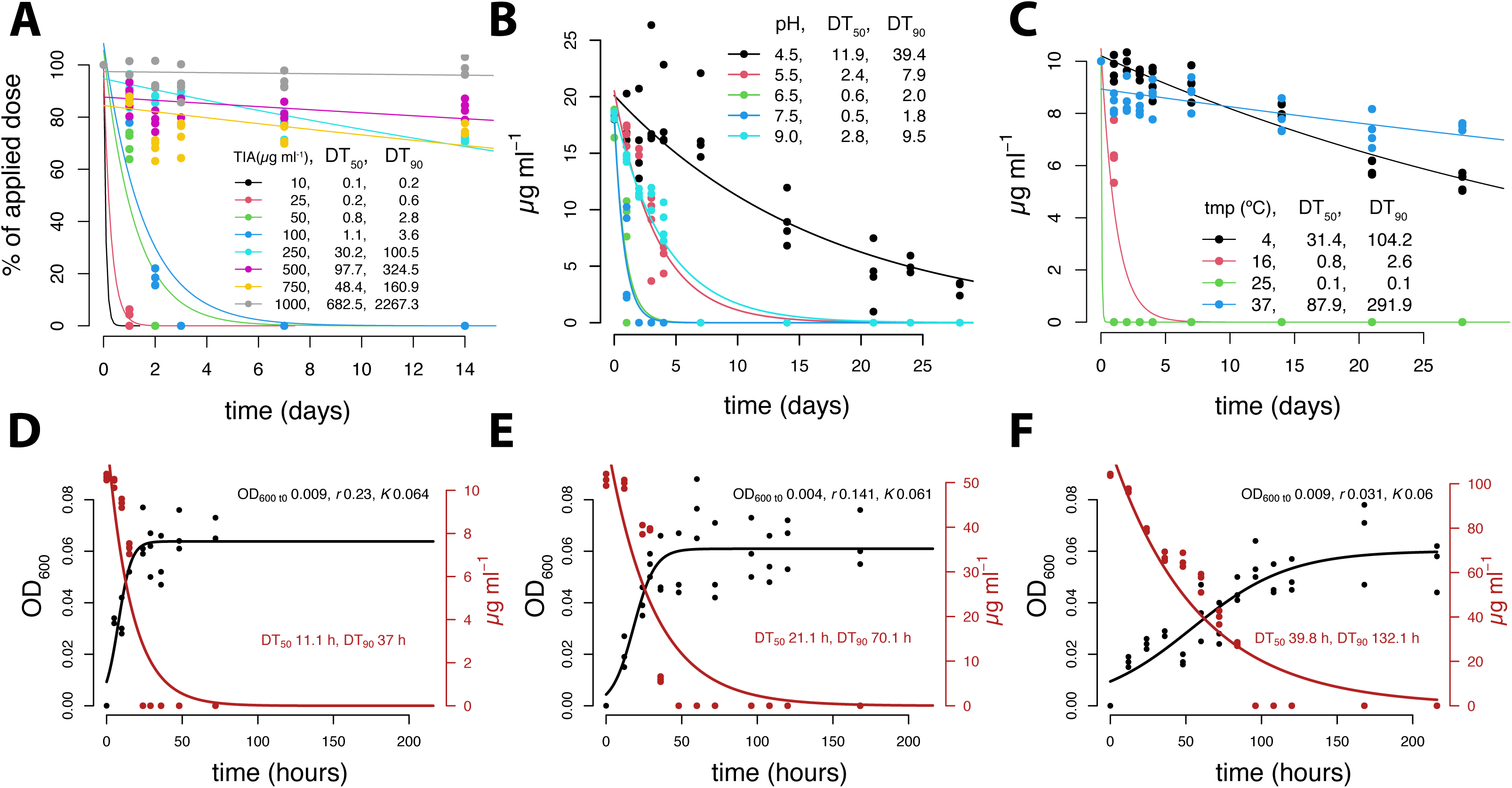
Characterization of the degrading capacity of the TIA-degrading bacterium. The degradation kinetic patterns of TIA at (A) different concentration levels: 10, 25, 50, 100, 250, 500, 750, and 1000 µg ml^-1^; (B) different pH conditions: 4.5, 5.5, 6.5, and 7.5 and (C) different temperatures: 4, 16, 37 °C. The degradation kinetic patters of TIA and the growth patterns of the TIA-degrading bacterium at three concentration levels of 10, 50 and 100 µg ml^-1^ are shown in D, E and F. The DT_50_ and DT_90_ values of TIA were calculated according to the single first order (SFO) model which showed the best fit to the data in all cases. The growth curve parameters of OD_600_ at time 0 h, the maximum growth rate (*r*) and the carrying capacity (*K*; or maximum OD_600_) were calculated according to the logistic model implemented by the growthcurver R package.

To verify that the isolate is using TIA as a growth substrate we monitored its growth in MSM in the presence of the different antibiotic concentrations: 10, 50 and 100 µg ml^-1^. TIA had DT_90_ values of 37, 70.1 and 132.1 h with respective inversely dose-dependent maximum growth rates (*r*) of 0.23, 0.141, and 0.031 (Figure 1D, E, F). All values and abiotic control data are provided in Figure S5.

### Genomic analysis

#### Genome assembly statistics, purity and phylogenomic analysis

The performed hybrid assembly resulted in a single contig of a total length of 5,168,335 bp without any gaps (Files S1, Table S1). Purity analysis with MiGA (36), suggested a high-quality assembly, with 100 % completeness and 0.9 % contamination (overall 95.5 % quality).

The Genome Taxonomy Database Toolkit (GTDB-Tk – v2.3.2) (37) analysis showed an average nucleotide identity (ANI) of 84.36 % over an alignment considering the 97.74 % of the isolate genome (Table S2), between the genome of our isolate and its closest relative *Sphingomonas laterariae* (Figure 2). This percentage is considered to be far below the 95 % ANI threshold above which strains of the same species reside (37, 38), suggesting that the isolate comprises a candidate novel species according to the current GTDB (r214), which was named *Candidatus* Sphingomonas perruchonii DBB INV C78.

**Figure 2.**
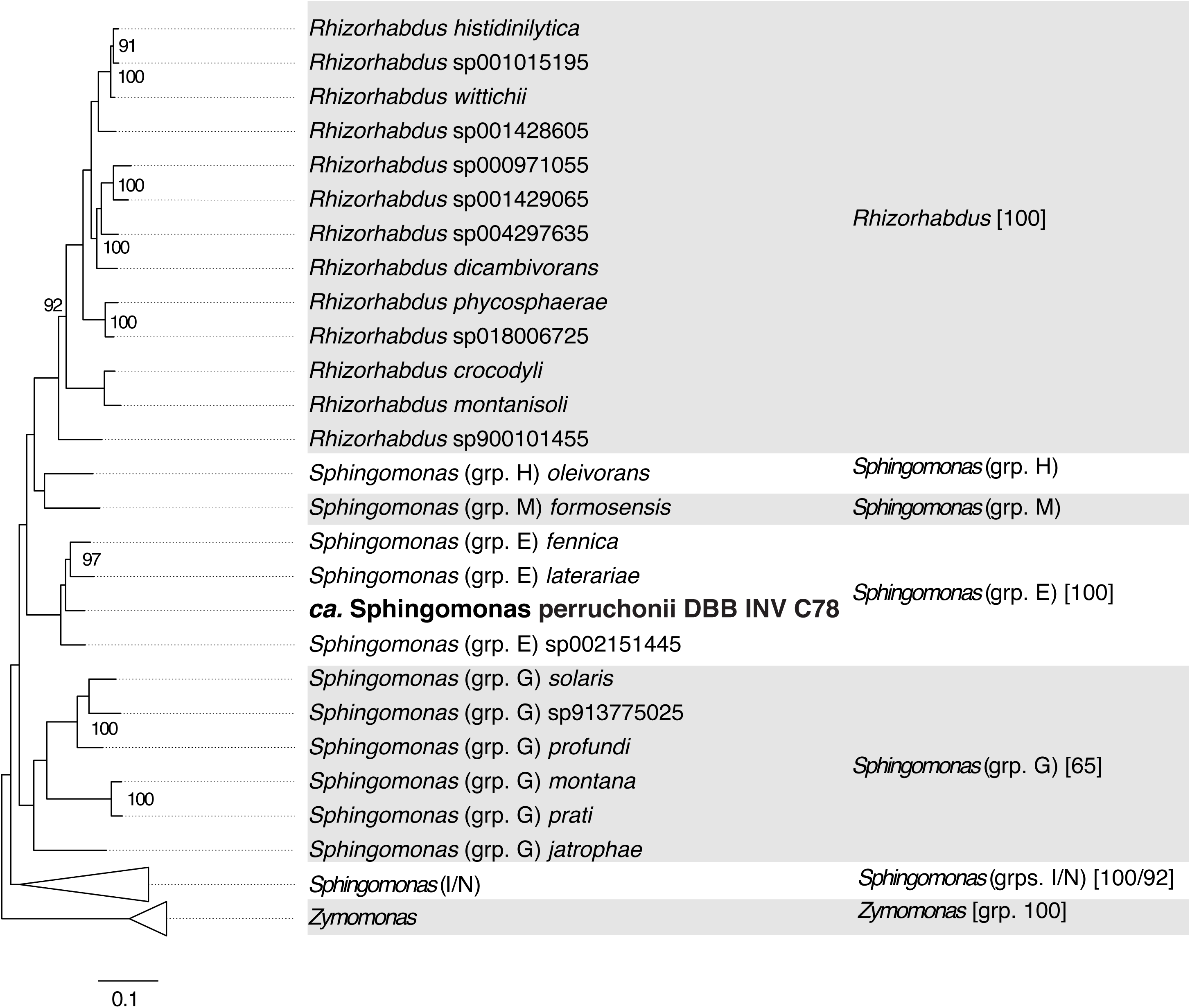
Phylogenomic closest-relatives subtree of the placement of the isolated microorganism aligned genomic loci, on the universal GTDB tree. The isolate is referred as *Ca*. Sphingomonas perruchonii DBB INV C78 given the low ANI value (84.36 %) to *Sphingomonas laterariae*, being the closest relative according to pairwise ANI comparisons (Table S2). Bootstrap values (where ≥ 75) are provided on the tree and, for species groups, in the square brackets of the rightmost column depicting these groups along with the background shading effect.

### Plasmid identification, insertion sequences prediction and resistome annotation

A locus spanning 7160 bp showed 84.75% identity with genes previously found to reside on plasmids according to PLASMe v1.1 (39), yet no hits were identified by the PlasmidFinder v2.1 pipeline (40). Most of these genes were associated with metal efflux pumps (cobalt-zinc-cadmium) or were annotated as hypothetical suggesting no presence of plasmid in *Ca.* Sphingomonas perruchonii (Figure 6B, Table S6).

ISEScan v1.7.2.2 (41) predicted the presence of 19 insertion sequences (ISs) classified into four major families (IS*3*, IS*21*, IS*66*, and IS*110*). ISCompare v1.0.7 (42) confirmed 15 of the ISEScan annotated ISs as mobilization events against the closest genome relative (*Sphingomonas laterariae* genome sequence GCF_900188165.1; Figure 6B, Tables S7-8, Figure S2).

Sixteen antimicrobial, antibiotic, and metal/metalloid resistance genes were identified by DFAST v1.2.10 (43) (Table S9). These were grouped into 6 metal/metalloid resistance (Cu transport/resistance (44) and As resistance genes), one disinfectant resistance gene (quaternary ammonium-resistance protein *SugE* (45)), a toxic anion resistance gene possibly linked to tellurite resistance, and six ARGs. These encompassed three multidrug efflux pumps, a Bcr/CflA family efflux pump homologous to bicyclomycin, chloramphenicol, florphenicol, and tetracycline resistance genes, a tetracycline efflux pump, a chloramphenicol resistance permease (RarD), an EmrB/QacA family drug resistance transporter conferring resistance to antibiotics, quaternary ammonium compounds and divalent cations (46, 47), and a glyoxalase/bleomycin dioxygenase. The CARD RGI v6.0.3 (48) analysis, although validated most DFAST ARG annotations, identified many more putative ARGs (395) of low probability (loose cutoff values), and two *adeF* ARGs coding for multidrug resistance efflux pumps (Tables S10-11).

Resistance to TIA has been previously linked with: (a) mutations at the peptidyl transferase loop near the peptidyl transferase centre (PTC) of the 23S rRNA at universally conserved nucleotide positions (49, 50); (b) a mutation at the 149^th^ amino acid (*E. coli* numbering) of the ribosomal protein L3 (uL3) interacting with the PTC (49, 50); (c) the presence of an ABC-F transporter (51). Ribosomal 23S rRNA gene BLAST analysis between *Escherichia* coli and our strain depicted the presence of a single point mutation out of the nine possible mutations previously linked with resistance phenotypes (Figure S6). Screening for the TIA-related uL3 mutation in our strain returned no hits (Figure S7). Finally, four putative ABC-F transporters were identified according to our CARD search (ACFB49_25800, ACFB49_34920, ACFB49_38340, ACFB49_44030; Tables S10-11). ACFB49_25800 was further investigated due to its interesting expression profile (described further below) and showed relative conservation against homologues with demonstrated contribution to resistance to TIA (Figure S7).

### Transcriptomic analysis

#### Transcriptomic profile of the analysed samples

Hypothesis testing showed a strong treatment effect on the transcriptional profiles (Figure S8), with the PERMANOVA model explaining 68.7% (*P* 0.001) of the total variance, partitioned at 36/64 % between treatment/time (*P* 0.0007/0). Treatment had a much stronger effect (*R^2^* 72%, *P* 0.0001) for a much better fitting overall model (*R^2^* 91%, *P* 0), when the 72-hour samples (30 hours after TIA degradation (where profile convergence between the two treatments was observed) were removed (Figure S8). Hence, only the 18- and 32-hour samples were considered in the identification of differentially expressed genes under TIA vs SUC treatments.

### TIA-induced differential gene expression

A total of 295 genes were identified as differentially expressed (DE) (*P* value 0.001 and a log_2_ fold change ≥ 2) (Figure 3A, Table S12). Out of these, 142 genes were significantly upregulated under the TIA treatment, and 28 of them had at least 64 times higher expression in the presence of TIA compared to their expression levels in the SUC treatment (Figure 3A, Table S13). These genes were mainly organized into two genomic regions, labelled “locus1” (parted by 6 genes and a total length of 4 Kbp) and “locus2” (parted by 64 genes and a total length of 78.5 Kbp) (Figure 3B, C and D and Table S14).

**Figure 3.**
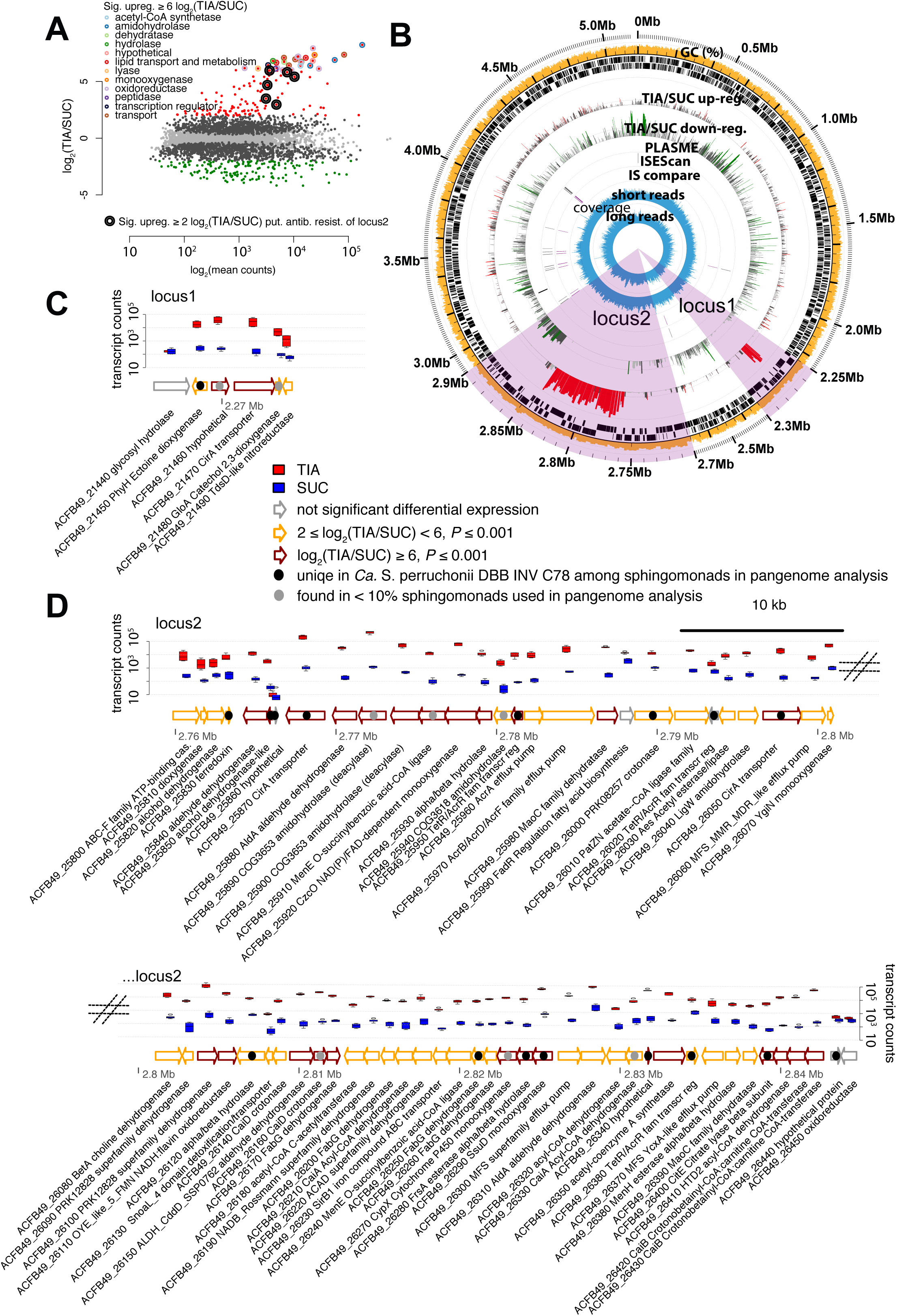
(A) Scatter plot of the transformed mean counts *vs* log_2_showing the differential expression between the two treatments (TIA vs SUC). Two hundred and ninety five genes were identified as differentially expressed (DE) for a *P* value and a log_2_(fold change) cutoff values of 0.001 and 2 respectively (red or green points for upregulated or downregulated genes respectively) with 28 of them being highly upregulated in the presence of TIA compared with SUC (log_2_ fold change value of 6). Circles of different colours show the broad functional categories of these 28 genes, while the double-lined black circle denotes the only antibiotic resistance gene (ARG) having significant 2-fold upregulation in TIA vs SUC. Light-grey points show the non-DE genes for both set criteria, while the dark grey points show genes that passed the P value criterion but not the log_2_ fold change cutoff criterion. (B) Circleator generated overview of the isolate genome with annotations showing (from inside-outwards): the long and short read coverage of the genomic areas; the insertion sequence locations as identified by IS compare and ISEScan; plasmid associated sequences identified by Plasme; TIA/SUC downregulated (green) and upregulated (red) genes; the gene representations (black bars) in either direction; the GC % content. Loci with highly upregulated genes under the TIA treatment (≥ 6 log_2_(TIA/SUC) were magnified x25 (locus1 and locus2). Finally, a more detailed overview of the genetic arrangement and gene expression in the TIA and SUC treatments (depicted with boxplots of the transcript counts at each treatment) is provided for locus1 (C) and locus2 (D).

Locus 1 comprised with two highly upregulated genes in the TIA treatment (log_2_ fold change ≥ 6) annotated as *tonB* receptor (ACFB49_21470) and a hypothetical protein (ACFB49_21460), and four genes which were also upregulated in the presence of TIA but to a lower extent (log_2_ fold change ≥ 2 and ≤6). These encoded a hydrolase (beta-glucosidase; ACFB49_21440) phytanoyl-CoA type dioxygenase (ACFB49_21450), an oxidoreductase (VOC family; ACFB49_21480), and an oxidoreductase (NAD(P)H-flavin; ACFB49_21490).

Locus 2 was composed of 64 genes functionally annotated as: oxidoreductases (26), hydrolases (10), lyases (6), ligases (4), transferases (2), regulatory proteins (5), transport/detoxification proteins (9), and hypothetical proteins (2). The group of oxidoreductases was dominated by dehydrogenases (20) and monooxygenases (4) while a dioxygenase (ACFB49_25810) and a ferredoxin (ACFB49_25820) were also noted as upregulated during TIA degradation. Amongst them, a cytochrome P450 monooxygenase (ACFB49_26270), a luciferase family monooxygenase (ACFB49_26290) and a FAD-dependent monooxygenase (ACFB49_25920) were highly upregulated in the TIA treatment (log_2_ fold change ≥ 6). Regarding hydrolases, we identified: (a) five amidohydrolases (ACFB49_25890, ACFB49_25900, ACFB49_25940, ACFB49_26100) which, all but one, (ACFB49_26040, 2≥log_2_ fold change ≤6) showed particularly high levels of upregulation (log_2_ fold change ≥ 6) in the TIA treatment; and (b) two alpha/beta hydrolyses (ACFB49_25930, ACFB49_26280) that encompass a broad group of enzymes involved in the hydrolysis of several organic pollutants including antibiotics (52).

Besides genes with putative role in the degradation of TIA (monoxygenases and (amido)hydrolases), the rest of the upregulated genes were involved in major cellular processes: (a) fatty acid metabolism like lyases (ACFB49_25980, ACFB49_26000, ACFB49_26140, ACFB49_26160, ACFB49_26390), ligases (ACFB49_25910, ACFB49_26240) and transferases (ACFB49_26420); (b) energy producing like the citric acid cycle (ACFB49_26400), pyruvate/propanoate and acetate metabolism (ACFB49_26010, ACFB49_26350); and (c) multiple others like an acetyl-CoA C-acetyltransferase (ACFB49_26180). We also noted a high upregulation of regulatory proteins linked to: (a) antibiotic resistance efflux pumps (ACFB49_25950, ACFB49_26360); (b) fatty acid biosynthesis (ACFB49_25990, ACFB49_26020); (c) inorganic ion transport (ACFB49_26230). Finally, we identified several transport/detoxification related proteins that were strongly upregulated in the presence of TIA like: (a) putative antibiotic efflux pump components (ACFB49_25960, ACFB49_25970, ACFB49_26060, ACFB49_26300, ACFB49_26370); (b) toxic compound exporting enzymes (ACFB49_26130); (c) inorganic ion transportation components (ACFB49_25870, ACFB49_26050); and (d) an ABC-F transporter, associated with ribosomal protection from ketolides, lincosamides, macrolides, oxazolidinones, phenicols, pleuromutilins, and streptogramins (ACFB49_25800).

Twenty-five of the genes in these two loci either lacked homologues or had homologues at less than 10 % of the genomes used in the *Sphingomonas* pangenome analysis (Figure 3C and D; Table S15). The analysis showed an open genus (*α* of 0.60; way below 1 that indicates a closed pangenome) according to H. Tettelin et al. (53) with a core genome of just 546 genes and a pangenome of 20,694 genes (Figure S9).

### Shotgun analysis of the TIA transformation products

We further looked for the formation of potential transformation products (TPs) during biotic and abiotic degradation of TIA. Several high molecular mass compounds were detected at early stages of the experiment in both inoculated and non-inoculated cultures (Figure 4; Table S16) including: (a) tiamulin sulfoxide (TP1, Sulfoxide TIA, mass 509,324) and oxidation products of the antibiotic like hydroxy tiamulin (TP2, OH-TIA, mass 509.324); (b) hydrolysis products like N-deethyl-tiamulin (TP4, N-deethyl-TIA, mass 465.291) and (c) and the hydroxy N-deethyl-tiamulin (TP5, OH-N-deethyl-TIA, mass 481.293) which was a product of oxidation and hydrolysis of TIA. All these TPs were degraded within the first 50 to 125 hours only in the inoculated cultures denoting a potential role as transient products in the microbial transformation of TIA. In contrast, we observed the formation and accumulation, only in the inoculated cultures, of two compounds (Figure 4): (a) a di-hydroxylated derivative of TIA in its tricyclic moiety (TP3, di-OH-TIA, mass 525.319); and (b) a hydrolysis product of low molecular mass, the 2-diethylamino-ethyl-thio acetic acid and isoforms (TP9-11, C_8_H_18_NO_2_S, mass 192.105). Our analysis identified three other potential TPs of TIA (TP6, C_28_H_46_NO_3_S, mass 476.309; TP7, C_15_H_27_NO_3_S, mass 301.170; TP8, C_15_H_26_NO_3_S, mass 300.162) that were formed in equivalent amounts and showing similar formation patterns in both inoculated and non-inoculated cultures denoting their production through abiotic processes (Figure S10).

**Figure 4.**
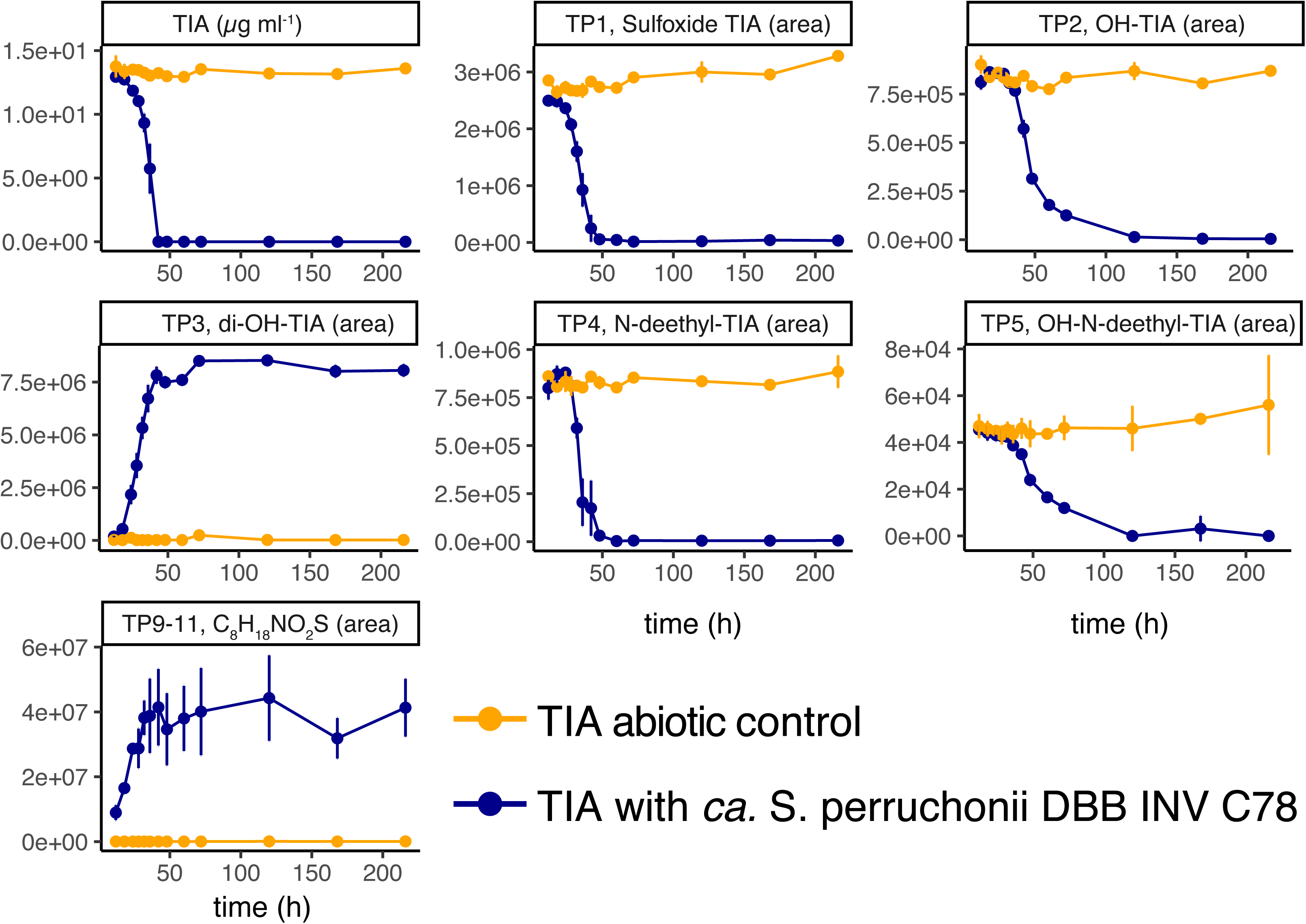
Temporal patterns of formation and decay of putative transformation products (TPs) of tiamulin (TIA) over time, with TIA measured in µg mL^-1^ whereas the rest products being measured in liquid chromatography peak areas, for the TIA abiotic controls and the TIA containing medium inoculated with the *Ca.* Sphingomonas perruchonii DBB INV C78.

## DISCUSSION

Microbial antibiotrophy is promising for a cost effective and environmentally friendly *in situ* reduction of antibiotic selective pressure. It is generally considered that exposure to antibiotics is necessary for the development of degradative traits within the microbial community. It is currently known that bacteria in natural habitats harbour both ARGs and genes associated with growth linked degradation of several antibiotics, including synthetic ones (23, 54, 55). Soil, in particular, owing to its enormous biodiversity and range of physical/chemical properties, is a promising source of such traits. Several studies (54, 56, 57) showed that soil bacteria can subsist on antibiotics as source of nutrients and energy, even when not previously exposed to it. In addition, spontaneous mutations can occur even in the absence of antibiotics (58).

In the present study, we relied on a soil that demonstrated accelerated dissipation of TIA (6) for to the successful isolation of our TIA-degrading bacterial strain. The specific soil did not have a history of TIA exposure, however its repeated treatment with TIA under laboratory conditions (34) has probably resulted in the selection of bacteria producing enzymes able to modify molecular sub-structures like those found in TIA, and eventually in their isolation.

Phylogenomic analysis suggested that the isolate was most closely associated with *Sphingomonas laterariae,* a hexachlorocyclohexane-degrading bacterium isolated from a contaminated dump site in India (59). However, the calculated ANI of 83.87% between the two close genomes suggests that the TIA-degrading isolate should be considered a candidate novel species which was named *Candidatus* Sphingomonas perruchonii DBB INV C78. *Sphingomonas* is known for the general ability to degrade refractory pollutants, in particular aromatic hydrocarbons, thanks to the assortment of unusual anabolic and catabolic pathways harbored (60–64).

*Ca.* Sphingomonas perruchonii was able to degrade and grow on 100 µg ml^-1^ TIA but failed to degrade higher concentration levels tested (≥ 250 µg ml^-1^). We speculate that at those high TIA concentrations, the biocidal activity of the antibiotic could not be counteracted by the antibiotic resistance mechanisms of the bacterium, thus hampering its degrading capacity.

This is further supported by the inverse association of the maximum growth rates of the isolate with the TIA nominal concentrations (Figure 1). *Ca.* S. perruchonii maintained its degrading capacity at a wide range of pH (4.5 to 9) and temperatures (16-25°C) suggesting, along with its capacity to degrade high TIA levels, a high potential for inclusion in bioremediation strategies of TIA – contaminated manures or soils (65).

We further employed a combination of genomic, transcriptomic and high-resolution untargeted LC-MS/MS analysis to (a) explore the potential mechanism of resistance to TIA and (b) disentangle the transformation pathway of TIA and the genetic elements involved. The complete genome of *Ca.* Sphingomonas perruchonii consisted of a single circular chromosome of around 5.1 Mbp with 4,903 coding sequences (CDS), which is within the range of genome sizes (2.8 to 6.5 Mb) and CDS number (1993 to 5951) reported for other *Sphingomonas* strains (60, 66).

Considering the capacity of our isolate to tolerate TIA, besides degrading it, we investigated its genome for mechanisms conferring resistance to antibiotics. Members of the *Sphingomonadaceae* can be generally considered a potential reservoir of antibiotic resistance, with *Sphingomonas* being the hotspot of antibiotic resistance traits (67). We initially sought for known mutations linked to the reduction of the affinity of TIA to the 23S rRNA peptidyl transferase center (PTC) both on the large ribosomal subunit (LSU) and the uL3 protein (49, 50), without much success (Figures S6,S7A). On the other hand, we identified a highly upregulated ABC-F transporter, without a transmembrane domain, whose function is to protect the LSU from TIA and possibly other antibiotics of similar modes of action (68). This transporter showed relative conservation of its activity-specific domains with other ABC-F transporters (Figure S7), previously linked with TIA resistance (51). However, this ABC-F transporter was only able to confer resistance to TIA levels between 0.25-8 µg ml^-1^ (51), way below the 100 µg ml^-1^ of TIA tested here. The CARD and DFAST search identified seven efflux pump associated genes that were overexpressed under the TIA treatment: three MFS, two TetR/AcrR and two Acr efflux pump transcriptional regulators (Table S14). Both the Acr and MFS efflux pumps are known to reside in multi-drug resistant bacteria (69), with the latter having extremely relaxed substrate specificity (70, 71). Our data suggest that resistance to TIA is potentially derived through a coordinated mechanism involving multiple modes of resistance like the protection of the antibiotic target by detoxification of its immediate environment (ABC-F transporter) and the detoxification of the cells in their entirety (ACR and MFS efflux pumps).

In the quest to identify candidate genes involved in the transformation of TIA, we noted a range of alpha/beta hydrolases, (amido)hydrolases and monoxygenases/dioxygenases known to play central roles in the biodegradation of organic pollutants (72), in line with the profile of an antibiotroph (19). Pangenome analysis showed relatively low dispersal of these genes among sphingomonads, highlighting the uniqueness of the isolate studied here regarding TIA degradation.

Non-target LC-MS/MS analysis identified several potential TPs of TIA, including both oxidation and hydrolysis derivatives (Figure 5, Table S2). Some of these TPs were formed only in the inoculated cultures validating their biological nature. Out of those TPs, only two, a mono-hydroxylated derivative of TIA in the tricyclic moiety and 2-diethylamino-ethyl-thio acetic acid (product of the cleavage of the aliphatic chain of TIA from the tricyclic moiety), were found to accumulate in the medium, while the rest were of transient nature. Based on these data we speculated that TIA undergoes successive oxidations in the tricyclic moiety with the hydroxy-derivative further hydrolysed to 2-diethylamino-ethyl-thio acetic acid or its isoforms, unlike the di-hydroxylated derivative which accumulates in the medium.

**Figure 5.**
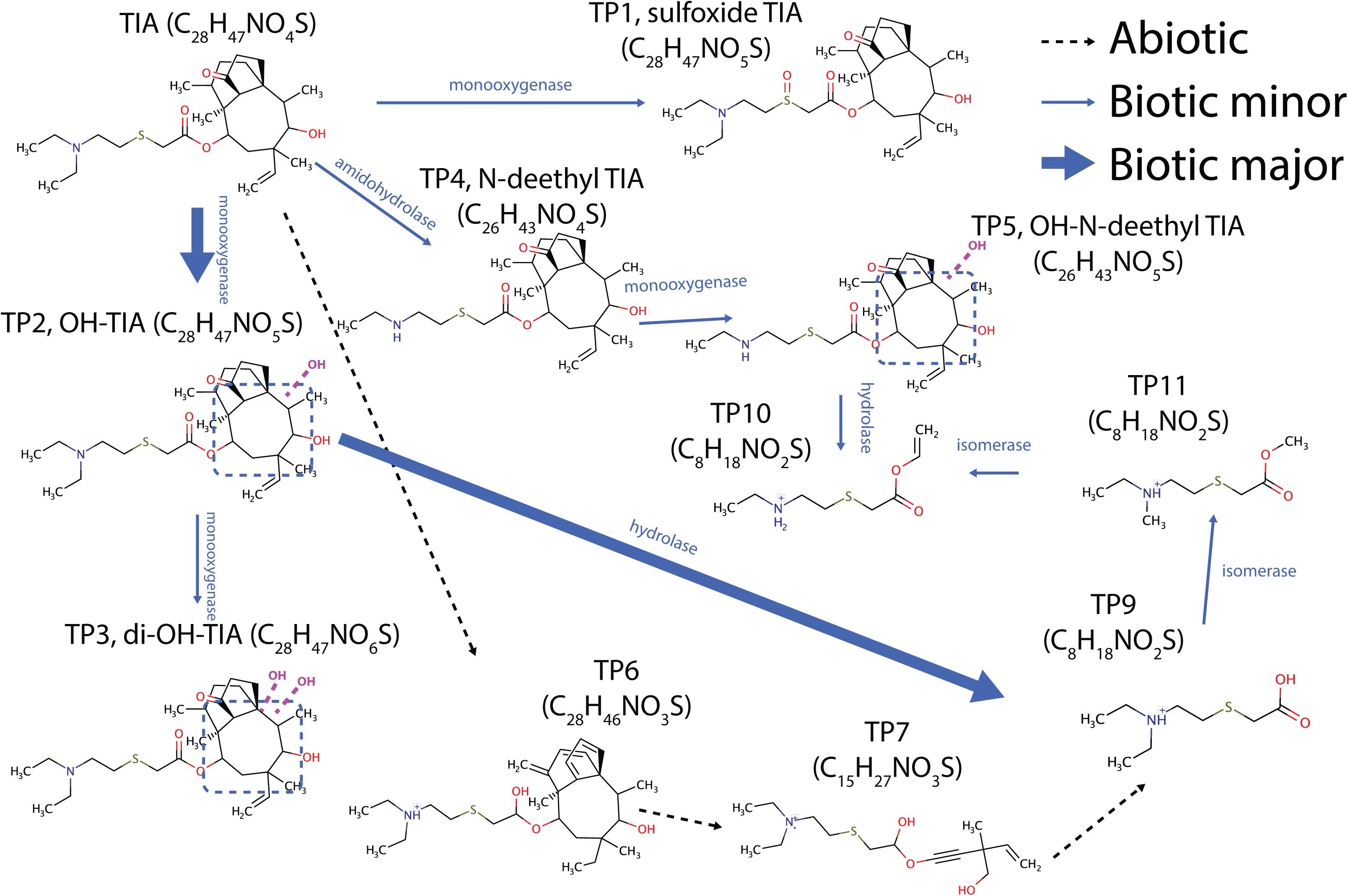
The proposed metabolic pathway for TIA transformation by the isolated bacterium. Dashed purple lines and blue-framed areas indicate uncertainty of the exact oxidation location.

When the results of the LC-MS/MS analysis and the transcriptomic analysis were coupled, we were able to identify suspect genes encoding for enzymes responsible for the different steps of the potential transformation pathway of TIA. Hence the highly upregulated monooxygenases (e.g. Cytochrome P450 or cyclohexanone FAD-dependent monooxygenase) or dioxygenases are probably involved in the hydroxylation of TIA towards the production of OH-TIA and di-OH-TIA. Modifications of the tricyclic core of the TIA molecule are considered challenging due to a limited number of functional groups (73). However, hydroxylation of the mutilin part (the ring system) by *Streptomyces griseus* at the 7- and 8-positions of the ring have been observed by R. L. Hanson et al. (74), probably due to the activity of a cytochrome P-450 monooxygenase-like enzyme. Moreover, hydroxylations in the ring system, together with S-oxidation and *N*-desethylation on the side chain, have been identified as the main modifications that TIA undergoes in its *in vivo* biotransformation in farm animal bodies (75). Hydroxylations introduce functional groups allowing for easier subsequent modification of the molecule: the insertion of hydroxyl functions on the ring, destabilize it making it more prone to further hydrolysis (61, 76, 77). Indeed, we noted several amidohydrolases and alpha/beta hydrolases residing in locus 2 that were highly upregulated during TIA degradation. These enzymes could be responsible for the hydrolysis of TIA to 2-diethylamino-ethyl-thio acetic acid and its isoforms, the latter could be also formed through the action of the highly up-regulated amidohydrolases. These low molecular mass TPs constitute the lateral chain of the TIA molecule, which replaces the hydroxyacetate group of pleuromutilin in the synthesis of TIA (78). The observed transformation pathway of TIA can be considered a detoxification process, since it leads to the accumulation of a final, to our knowledge, non-toxic compound, which constitutes a desirable trait for future applications in bioaugmentation strategies. Several previous studies have reported microbial transformation processes for antibiotics that lead to the formation of final products that exert the same or even higher toxicity compared to their parent chemical (79–82).

Based on the above, a putative transformation pathway for TIA is proposed (Figure 5). TIA is subjected to successive oxidation of its tricycling ring which leads to the formation of mono-oxygenated (TP2) and dioxygenated derivatives (TP3). The latter is not further degraded and accumulated in the culture, unlike the former which is further transformed most probably to 2-diethylamino-ethyl-thio acetic acid (or its isoforms) (TP9, 10 or 11) (Figure 5). Other minor biotic transformation paths like (i) the sulfoxidation of TIA and the (ii) desethylation and subsequent oxidation of the tricyclic moiety of the desethylated TIA were also active.

### Concluding remarks

We report here the isolation of the first bacterium able to degrade and grow on TIA, a very commonly used veterinary antibiotic. Phylogenomic analysis revealed that the isolated strain could be classified as a new species of Sphingomonas, baptized *Candidatus* S. perruchonii DBB INV C78. Genomic analysis revealed typical features of antibiotrophism like (i) a multifaceted toolbox for resistance to antibiotics and (ii) a rich arsenal of genes with potential role in the catabolism of xenobiotics, most of them being highly upregulated during degradation of TIA. Combination of transcriptomic and non-target metabolomic analysis, led us to define a possible transformation pathway of TIA and pointed to genes encoding enzymes that might be involved in the main steps of this pathway. Our results suggest that the isolated bacterium could be a good candidate for future applications aimed at the bioaugmentation of contaminated manures and soils. Further tests will focus on the verification of the role of the highlighted monooxygenases and (amido)hydrolases in the transformation of TIA through targeted mutagenesis, heterologous expression and *in vitro* functional testing.

## MATERIALS AND METHODS

### Chemicals and growth media

Tiamulin fumarate (Biosynth® Carbosynth, Staad, Switzerland – purity >97 %) was used in all assays. The selective medium used for the isolation and routine cultivation of the TIA-degrading bacterial strain was a mineral salt medium plus casamino acids (0.15 g l^-1^) which was either supplemented (MSMN) or not supplemented (MSM) with nitrogen (35). TIA was added in MSMN or MSM as the sole C, or C and N, source respectively, at a concentration of 10 µg ml^−1^, unless otherwise stated, using a filter-sterilized DMSO stock solution (15000 µg ml^−1^). The Luria Bertani medium was also used for purity testing of the isolates.

Corresponding solid media of MSM and MSMN were obtained by adding 15 g agar l^-1^ prior autoclaving. For purifying the TIA degrading isolate, several antibiotics were added to the growth medium at various concentrations: chloramphenicol (15 µg ml^-1^), kanamycin (30 µg ml^-1^), tetracycline (60 µg ml^-1^), rifampicin (0.5, 1, and 5 µg ml^-1^), gentamycin (1, 2, and 10 µg ml^-1^), and ampicillin (5, 10, and 20 µg ml^-1^). All antibiotics were purchased by Sigma-Aldrich at ≥ 99% purity (St. Louis, MO, USA).

### Tiamulin residue analysis

TIA was extracted from liquid media by mixing an aliquot of the liquid culture with methanol, vortexing for 30 sec and centrifuging for 3 min at max speed. The clear supernatant was collected and stored at −20°C until downstream analysis. Samples were analysed in a Shimadzu HPLC-PDA system equipped with a CNH Athena RP C18-150 mm column (CNW Technologies, Düsseldorf, Germany) as described by C. Perruchon et al. (6).

### Isolation of a TIA-degrading microorganism through enrichment

A soil with demonstrated accelerated TIA dissipation (6) was used as a source for the isolation of TIA-degrading microorganisms. The soil was subjected to three repeated applications of TIA (2.5 mg kg^-1^) at 50-day intervals, aiming to stimulate the TIA-degrading microbial community. An enrichment liquid culture strategy in selective media with TIA as a substrate was employed to isolate TIA – degrading microorganisms as previously described (83, 84). Flasks of MSM/MSMN supplemented with TIA (10 µg ml^-1^) were inoculated with the acclimated soil described above (2.5% w/v) and the cultures were incubated shaking in the dark at 25 °C. Inoculated and non-inoculated (control) triplicates were prepared. TIA degradation was monitored by HPLC and when more than 50% degradation occurred an aliquot of the degrading culture (5% v/v) was refreshed in fresh medium. After four enrichment cycles, a 10-fold dilution series of the degrading culture were prepared and spread on MSM/MSMN agar plates supplemented with TIA (10 µg ml^-1^). After incubation at 25°C for 4-5 days, single growing colonies were selected, inoculated in the respective liquid medium, and the TIA concentration was measured by HPLC. Cultures exhibiting > 50 % TIA dissipation in ≤ 7 days were considered positive and plated on LB+TIA agar plates to verify purity. When not pure, the spreading-selecting procedure was repeated. Pure TIA degraders, which were then used in downstream analysis, were obtained only after the addition to the media of different antibiotics at different concentrations.

### Characterization of the degrading capacity of the isolate

The tolerance and degrading capacity of the isolate at a broad range of TIA concentrations (25, 50, 100, 250, 750, and 1000 µg ml^-1^) was assessed in the MSM medium. TIA degradation was monitored at regular intervals post inoculation, while detailed analysis of the growth kinetics of the isolate on MSM + TIA was performed for the highest concentration level where the growth of the isolate was not inhibited. In that case, TIA degradation and bacterial growth were determined by HPLC and OD_600_ measurement respectively.

We further tested the optimal pH and temperature conditions for TIA degradation by the isolated microorganism. Hence, the degradation of 10 µg ml^-1^ of TIA was tested in MSM at pH values of 4.5, 5.5, 6.5. 7.5, and 9. Regarding temperatures, the degrading capacity of the bacterium in MSM + 10 µg ml^-1^ TIA were tested at 4, 16, 25, and 37°C. In all the above experiments, abiotic controls were retained and triplicates for each treatment were prepared. Degradation kinetics and microbial growth curve analysis were performed with the mkin v1.1.1 (85) and the growthcurver v0.3.1 (86) packages of the R software v4.1.3 (87).

### Genomic analysis

DNA extraction and sequencing of the near full length 16S rRNA gene, was employed for an initial assessment of the identity of the isolated strain as described previously (83).

Following, the genomic DNA extracts were sequenced at an iSeq instrument generating 150 bp paired-end reads (Illumina, San Diego, CA, USA), and with PacBio RSII generating hifi long reads as described in the supporting information. Unicycler v0.4.8 (88) was used for a hybrid genome assembly and taxonomic classification was performed with the Genome Taxonomy Database Toolkit (GTDB-Tk – v2.3.2) (37). Genome assembly purity was tested with the MiGA (36), automated annotation was performed with the DFAST v1.2.10 (43) and the RASTserver API (89). Deeper investigation on insertion sequences was performed with ISEScan v1.7.2.2 (41) and ISCompare v1.0.7 (42), plasmid search was performed with PLASMe PLASMe v1.1 (39) and PlasmidFinder v2.1 (40), and ARGs were further annotated with the CARD RGI v6.0.3 (48). Further manual curation was performed for genes and sequences of interest (e.g. insertion sequences and antibiotic resistance genes) as described in the supporting information. Finally, pangenome analysis was performed with the micropan v2.1 (90) and assisted by the microseq v2.1.6 R packages (91) for the estimation of the distribution of our own genome homologous genes among sphingomonads.

### Transcriptomic analysis

Triplicate cultures of the TIA-degrading isolate in MSMN were prepared and supplemented either with 50 µg ml^-1^ of TIA or with 83.5 µg ml^-1^ succinate (SUC), aiming to obtain the same carbon content in both treatments. Triplicate abiotic controls (TIA without inoculation) were also included. At times of 0, 12, 18, 24, 28, 32, 36, 42, 48, 60, 72, 219, 168 and 216 hours after inoculation, TIA degradation was determined via HPLC and bacterial growth was monitored via OD_600_ measurement. RNA was extracted from the liquid cultures at 18, 32, 72 hours post inoculation and directional mRNA sequencing libraries were prepared after rRNA removal using the QIAseqFastSelect 5s/16s/23s rRNA removal kit (Qiagen, Hilden, Germany), with the NEB Ultra II directional kit (New England Biolabs, Ipswich, Massachusetts, United States). The libraries were then sequenced at a NovaSeq instrument with the SP sequencing kit, generating 250bp paired-end reads (Illumina, San Diego, CA, USA). Generated sequences were quality controlled, and were mapped onto the isolate genome with STAR v2.7.7a (92), while the read counts per gene were calculated with htseq v0.11.2 (93). The gene expression tables were analysed for multivariate sample profiling and associated hypothesis testing as detailed in the supporting information.

### Shotgun untargeted analysis of the transformation products of tiamulin

Mirroring the RNAseq samples in the above experiment, samples were retrieved at 0, 12, 18, 24, 28, 32, 36, 42, 48, 60, 72, 120, 168 and 216 days from inoculated and non-inoculated cultures containing TIA for shotgun untargeted analysis of potential TPs of TIA. Aliquots of the microbial cultures were collected and mixed in a 50:50 ratio with methanol. After 30 sec vortexing, samples were centrifuged 1min/max speed/RT and further filtered through 0.22-μm PTFE syringe filters. LC-QTOF-MS/MS based identification of TIA and tentative candidate transformation products is detailed in the SI following the basic setup of our previous study (94).

The retrieved mass spectra were annotated with a database based on putative metabolite formulae of previous abiotic degradation studies (75), and formulae prepared *in silico*, as described previously (95). Briefly, the mass spectra of TIA were retrieved from the Wiley SpectraBase^TM^ (https://spectrabase.com, last visited at 1/6/2023) and peak intensities retrieved through electron ionization (EI) were extracted from the portable network graphics (PNG) formatted peak image with the Plot Digitizer software (https://plotdigitizer.com, last visited at 1/6/2022). Following, the CFM-ID v3.0 suite (96) was used for (a) fragment identification of the SpectraBase-retrieved Tiamulin spectra, or (b) for SMILES-based *in silico* spectral prediction. The rcdk v3.7.0 package (97) of the R software environment was used for chemical formula retrieval and structure plotting.

## Data availability

The genome and transcriptome sequencing data were submitted at the Sequencing Read Archive (SRA) and the Gene Expression Omnibus (GEO) of the National Center for Biotechnology Information (NCBI) and are publicly accessible under the Bioproject accession number PRJNA1168906.

## Acknolwedgments

The research project was financially supported by the Hellenic Foundation for Research and Innovation (H.F.R.I.) under the “2nd Call for H.F.R.I. Research Projects to support Post-Doctoral Researchers” (Project Number: 01183). The work performed by Niki Tagkalidou was further financially supported by the Postgraduate Program “Biotechnology – Quality assessment in Nutrition and the Environment”, Department of Biochemistry & Biotechnology, University of Thessaly (Project No. 5864).

